# Impulse initiation in engrafted pluripotent stem cell-derived cardiomyocytes can stimulate the recipient heart

**DOI:** 10.1101/2023.11.12.566756

**Authors:** Tim Stüdemann, Barbora Schwarzova, Till Schneidewind, Birgit Geertz, Constantin von Bibra, Marie Nehring, Judith Rössinger, J. Simon Wiegert, Thomas Eschenhagen, Florian Weinberger

**Affiliations:** Department of Experimental Pharmacology and Toxicology, University Medical Center Hamburg-Eppendorf, Germany; German Centre for Cardiovascular Research (DZHK), partner site Hamburg/Kiel/Lübeck, Germany; Research Group Synaptic Wiring and Information Processing, Center for Molecular Neurobiology Hamburg, University Medical Center Hamburg-Eppendorf, Hamburg, Germany; Department of Neurophysiology, MCTN, Medical Faculty Mannheim, Heidelberg University, Mannheim, Germany; Cardiovascular Regeneration Program, Centro Nacional de Investigaciones Cardiovasculares (CNIC), 28029 Madrid, Spain

## Abstract

Transplantation of pluripotent stem cell-derived cardiomyocytes is a novel promising cell-based therapeutic approach for patients with heart failure. However, engraftment arrhythmias are a predictable life-threatening complication and represent a major hurdle for clinical translation. Catheter-based electrophysiological analysis suggested that the ventricular arrhythmias were caused by an automaticity of the transplanted cells, but whether impulse generation by transplanted cardiomyocytes can propagate to the host myocardium and override the recipient rhythm has not been directly assessed experimentally. We used optogenetics to specifically activate engrafted cardiomyocytes, which resulted in impulse generation in the engrafted cardiomyocytes and stimulated the recipient heart (4/9 hearts). Thus, our study shows that transplanted cardiomyocytes can electrically couple to the host myocardium and stimulate the recipient heart, providing experimental evidence that cardiomyocyte automaticity can serve as a trigger for ventricular arrhythmias.

## Main

Transplantation of pluripotent stem cell-derived cardiomyocytes is a regenerative therapeutic strategy that holds great promise for patients with heart failure^1^. First clinical trials have started, and several others are currently in the planning stage. However, ventricular arrhythmias after cardiomyocyte transplantation (so-called engraftment arrhythmias) are the main hurdle for clinical translation. Engraftment arrhythmia is a common, potentially life-threatening complication that mainly occurs within the first three weeks after the transplantation^2–4^. Catheter-based electrophysiological studies in primates and pigs have provided evidence that it is caused by automaticity of the immature transplanted cells^4,5^. Here, we wanted to investigate whether impulse generation in the engrafted cells can propagate to the injured heart and override the recipient heart rhythm. We used optogenetics to specifically activate engrafted cardiomyocytes and show that impulse generation in the engrafted cardiomyocytes can stimulate the recipient injured heart. Hence, we provide evidence that cardiomyocyte automaticity can serve as a trigger for ventricular arrhythmias.

We transplanted induced pluripotent stem cell-derived cardiomyocytes which expressed the optogenetic actuator BiPOLES (that can be used to activate cardiomyocytes with blue and red light)^6,7^. This experimental set-up allowed us to specifically activate the engrafted cardiomyocytes with pulsed photostimulation. BiPOLES-iPSCs were differentiated to cardiomyocytes (average troponin T positivity: 93±4%) that were then injected transepicardially in a guinea pig injury model (30x10^6^ cardiomyocytes per animal, n=9). 56-days post-transplantation, the hearts showed transmural scarring (scar area was 27±3% of the left-ventricle, Figure 1A and B). Engrafted human cardiomyocytes (graft size: 17±7% of the scar, Figure 1 A and B) reached into the host myocardium and formed direct graft-host interactions (Figure 1C). Whereas connexin 43 was circumferentially localized, indicating that the engrafted cardiomyocytes were not fully matured, N-cadherin showed signs of polarization and formed intercalated-disc like structures (Figure 1D and E). Cardiomyocytes demonstrated advanced sarcomere structure and mainly expressed the ventricular (mature) isoform of the myosin light chain (93±2%, Figure 1F). A beginning switch from the fetal slow skeletal to the cardiac troponin I isoform (cardiac troponin I expression: 28±2%; Figure 1G) further indicated maturation. Cell cycle activity was still ongoing (8±2% Ki67 positive cardiomyocyte nuclei, Figure 1H).

**Figure 1:**
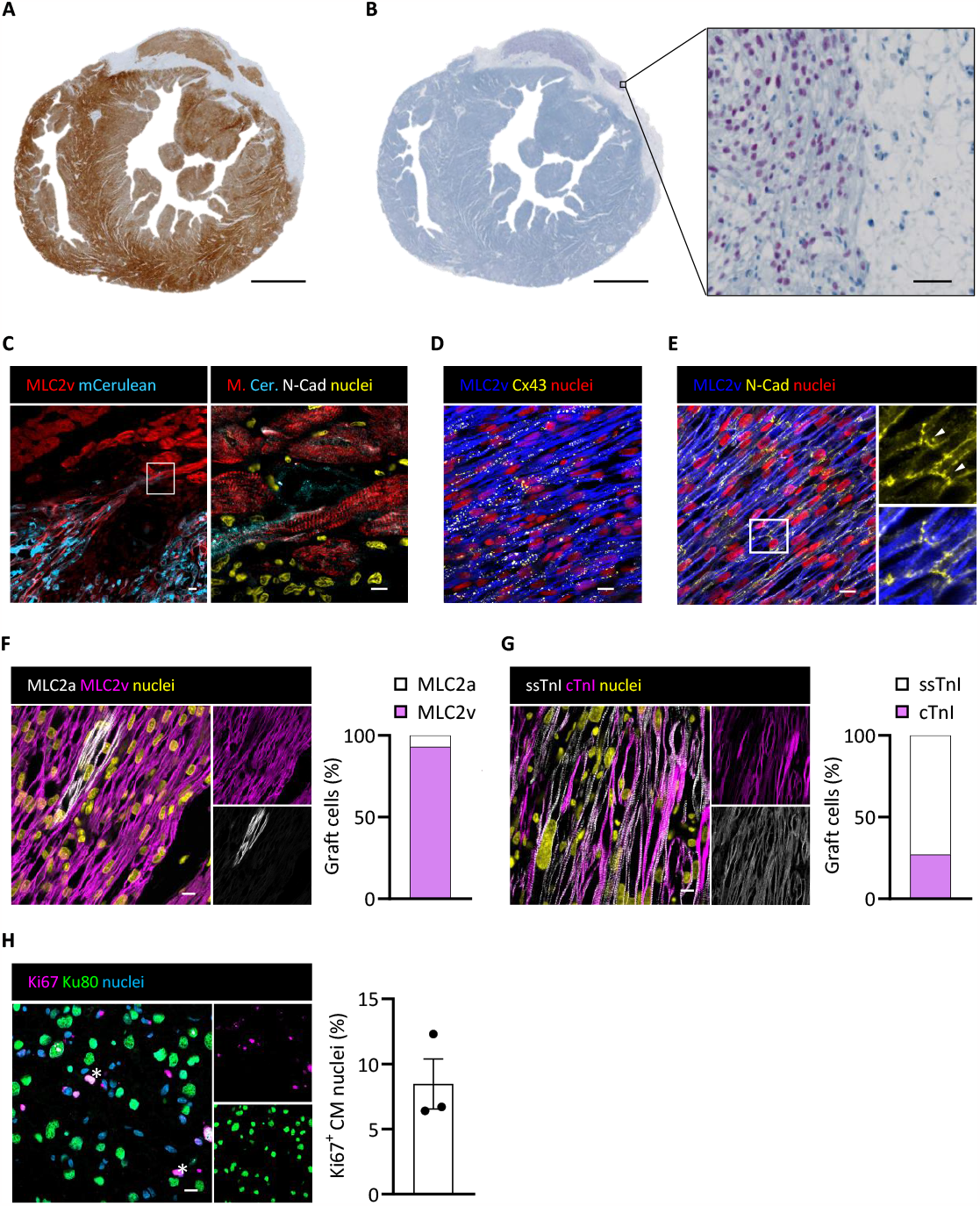
BiPOLES cardiomyocyte transplantation partially remuscularized the injured heart. **A-B**) Short axis sections of a guinea pig heart eight weeks after BiPOLES-cardiomyocyte transplantation stained for dystrophin (**A**) and human Ku80 (**B**). Inset in B is shown in higher magnification on the right. **C**) Graft-host interaction in lower and higher magnification. **D)** and **E)** High magnification images showing (**D**) connexin43 and (**E**) N-cadherin expression in the human grafts. Higher magnification depicts the inset to visualize an intercalated disc like structure. **F)** Staining for myosin light chain isoform expression. **G**) Staining for troponin I isoform expression. Quantification for E and F was based on pixel area per image. 4 images per heart from three different hearts were used for quantification. **H**) Analysis of cell cycle activity. Each data point represents one heart. Four images per heart were used for quantification. Asterisk mark human nuclei in the cell cycle. Scale bars: 2 mm in A and B. 100 μm in B (high magnification) and 10 μm in C-G.

Ex-vivo Langendorff-perfusion was used to assess electrical coupling. For this, the aorta was cannulated. A balloon catheter was used to measure left ventricular isovolumetric pressure and two ECG electrodes were placed on the surface of the heart (Figure 2A). Pulsed photostimulation (blue [470 nm] or red [635 nm] light, pulse duration 40-100 milliseconds) was applied to the site of injury at the anterior wall of spontaneously beating hearts (average beating frequency 181±15 bpm). The initial photostimulation frequency was adjusted to be higher than the spontaneous beating frequency (spontaneous beating frequency +0.25 to 1 Hz). Pulsed photostimulation resulted in ectopic pacemaking in 4 out of 9 recipient hearts (Figure 2 B -D). We then performed a stepwise increase in stimulation frequency. All hearts that showed electrical coupling could be paced up to 5 Hz, and the maximum pacing frequency was 10 Hz (Figure 2D). There was no difference between red and blue light photostimulation with respect to capture or maximum pacing frequency (which is in line with the in vitro characterization of BiPOLES in cardiomyocytes^7^). Adenosine (0.3 mg) was applied to inhibit atrio-ventricular conduction and to induce transient ventricular asystole. Photostimulation was initiated after the onset of ventricular asystole. Again, pulsed photostimulation resulted in optogenetic impulse generation that initiated a ventricular rhythm in 4 out of 9 hearts (Figure 2E and F). Comparing electrophysiological results to histological analysis revealed that cardiomyocytes were not engrafted in 4 of the 5 hearts that failed to respond to photostimulation. Consequently, 4 of the 5 hearts with engrafted cardiomyocytes exhibited ectopic pacemaking with pulsed photostimulation (Figure 2G). While there is evidence that impulse propagation can also occur via fibroblasts^8,9^ histology revealed structural coupling between engrafted cardiomyocytes and host myocardium in our study.

**Figure 2:**
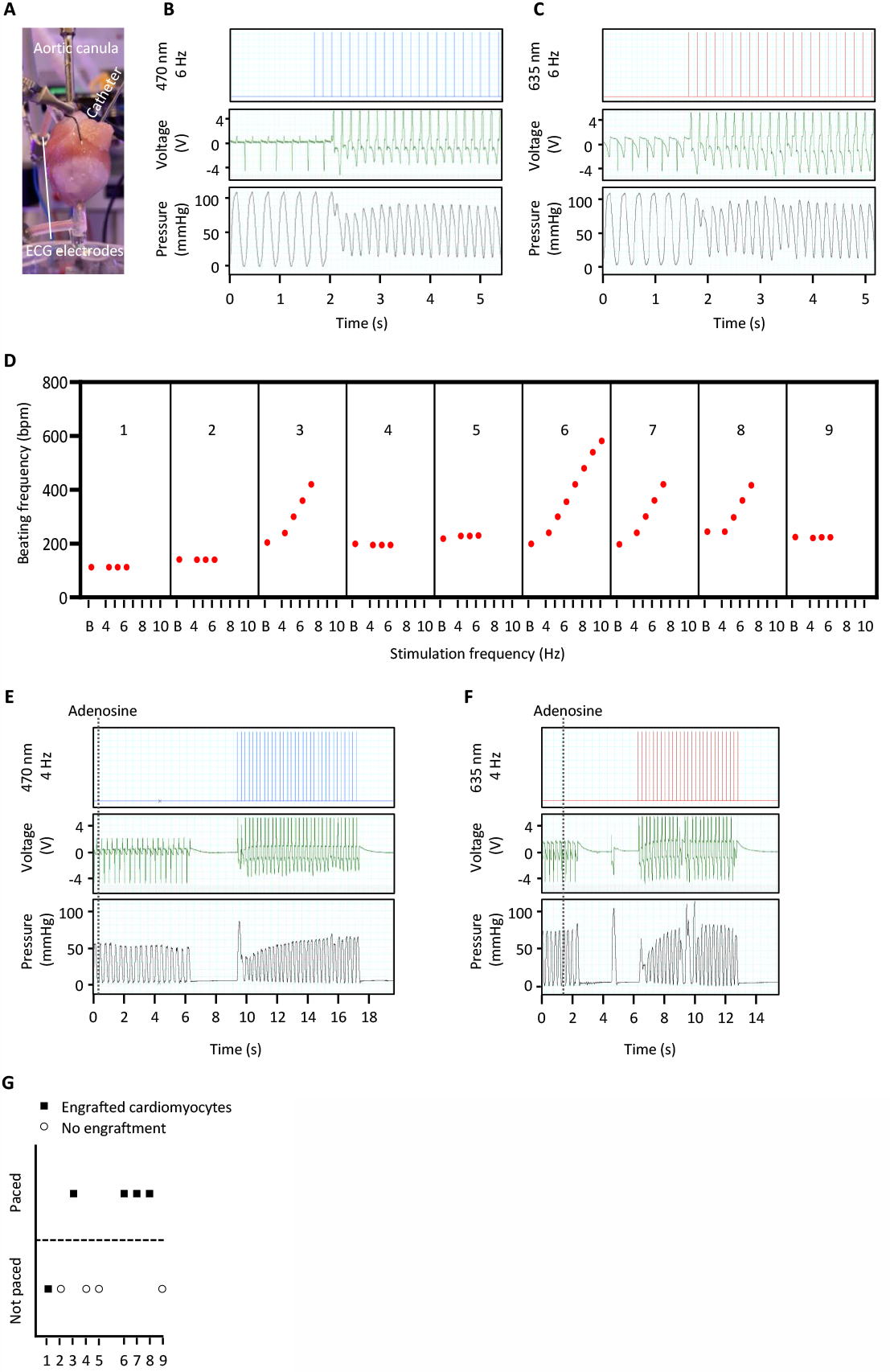
Impulse generation in engrafted cardiomyocytes stimulate the recipient heart. **A**) Photography of the experimental set-up. **B** and **C**) Original recording of an ECG (middle row) and left-ventricular pressure (lower row) under baseline condition and upon photostimulation with (B) blue light (470 nm, 6 Hz, upper row) and (C) red light (635 nm, 6 Hz, upper row). **D**) Quantification of pacing capabilities with red light. Nine individual hearts are shown. **E** and **F**) Original ECG and left-ventricular pressure recording while adenosine was applied via the coronaries. Pulsed photostimulation with (E) blue light (470 nm, 4 Hz) and (F) red light (635 nm, 4 Hz). was initiated after ventricular asystole was established. **G**) Comparison between coupling and cardiomyocyte engraftment.

Previous studies with genetically encoded calcium and voltage indicators provided evidence that engrafted cardiomyocytes can be stimulated by the recipient myocardium and beat in synchrony with host myocardium^10,11^. We used a contrary approach and stimulated the engrafted cardiomyocytes. By employing optogenetics, our study provides evidence that i) transplanted human cardiomyocytes electrically couple to the host myocardium and that ii) impulse initiation in the engrafted cardiomyocytes can stimulate the injured recipient heart. Thus, it supports the hypothesis that automaticity of the engrafted cardiomyocytes can trigger ventricular arrhythmias.

## Methods

### Human iPSC culture and cardiac differentiation

Generation of the BiPOLES iPSC line was recently described^7^. iPSCs were cultured in FTDA medium on Geltrex-coated cell culture vessels. Formation of embryoid bodies was performed in spinner flasks. Embryoid body formation was followed by differentiation with the sequential administration of growth factor- and small-molecule-based cocktails to induce mesodermal progenitors, cardiac progenitors, and cardiomyocytes. Cardiomyocytes were dissociated with collagenase. The protocol in detail has been described previously^12^. Cardiomyocytes underwent heat shock 24 hours before dissociation as previously described^13^ and were cryopreserved.

### Flow cytometry

Single-cell suspensions of iPSC-derived cardiomyocytes were fixated in Histofix (Roth A146.3) for 20 min at 4 °C and stained in a buffer, containing 5% FBS, 0.5% Saponin, 0.05% sodium azide in PBS. Antibodies are listed in Table 1. Samples were analyzed with a BD FACSCanto II Flow Cytometer and the BD FACSDiva Software 6.0 or BD FlowJo V10.

**Table 1.** Primary antibodies for flow cytometry.

| Antigen | Supplier | Titer |
| --- | --- | --- |
| Cardiac troponin T (APC) | Miltenyi Biotec, clone REA400, Cat. 130-120-403 | 1:50 |
| Isotype cardiac troponin T | Miltenyi Biotec, clone REA293, Cat. 130-120-709 | 1:50 |

### Animal care and experimental protocol approval

The investigation conforms to the guide for the care and use of laboratory animals published by the NIH (Publication No. 85-23, revised 1985) and was approved by the local authorities Behörde für Gesundheit und Verbraucherschutz, Freie und Hansestadt Hamburg: N098/2019 and N123/021).

### Injury model and cardiomyocyte transplantation

Cryoinjury of the left ventricular wall was induced in female guinea pigs (∼300 g, ∼8 weeks of age, Envigo) as previously described by our group^14,15^. Cardiomyocyte transplantation was performed seven days after the injury. During this procedure 30x10^6^ cardiomyocytes, resuspended in pro-survival cocktail^4,16^ (Matrigel (∼50% v/v), cyclosporine A (200 nM), pinacidil (50 μM), IGF-1 (100 ng/ml) and Bcl-X_L_ BH4 (50nM), total volume: 150 μl) were injected into three separate injection sites targeting central lesion and the flanking lateral border zones. Immunosuppression was performed with cyclosporine beginning three days prior to transplantation (7.5 mg/kg body weight/day for the first three postoperative days and 5 mg/body weight/day for the following 53 days) and methylprednisolone (2 mg/kg body weight/day).

### Histology

Hearts were sectioned into four to five slices after fixation. Serial paraffin sections were acquired from each slice and used for histological analysis. Antigen retrieval and antibody dilution combinations used are summarized in Table 2. The primary antibody was either visualized with the multimer-technology based UltraView Universal DAB Detection kit (Ventana® BenchMark® XT; Roche) or a fluorochrome-labeled secondary antibody (Alexa-conjugated, Thermofisher). Confocal images were acquired with an LSM 800 (Zeiss). For morphometry, images of dystrophin-stained sections were acquired with a Hamamatsu Nanozoomer whole slide scanner and viewed with NDP software (NDP.view 2.6.13). Graft size was measured in dystrophin-stained short-axis sections and expressed as a percentage of the scar area measured in the same section with the NPD2.view-software.

**Table 2.**
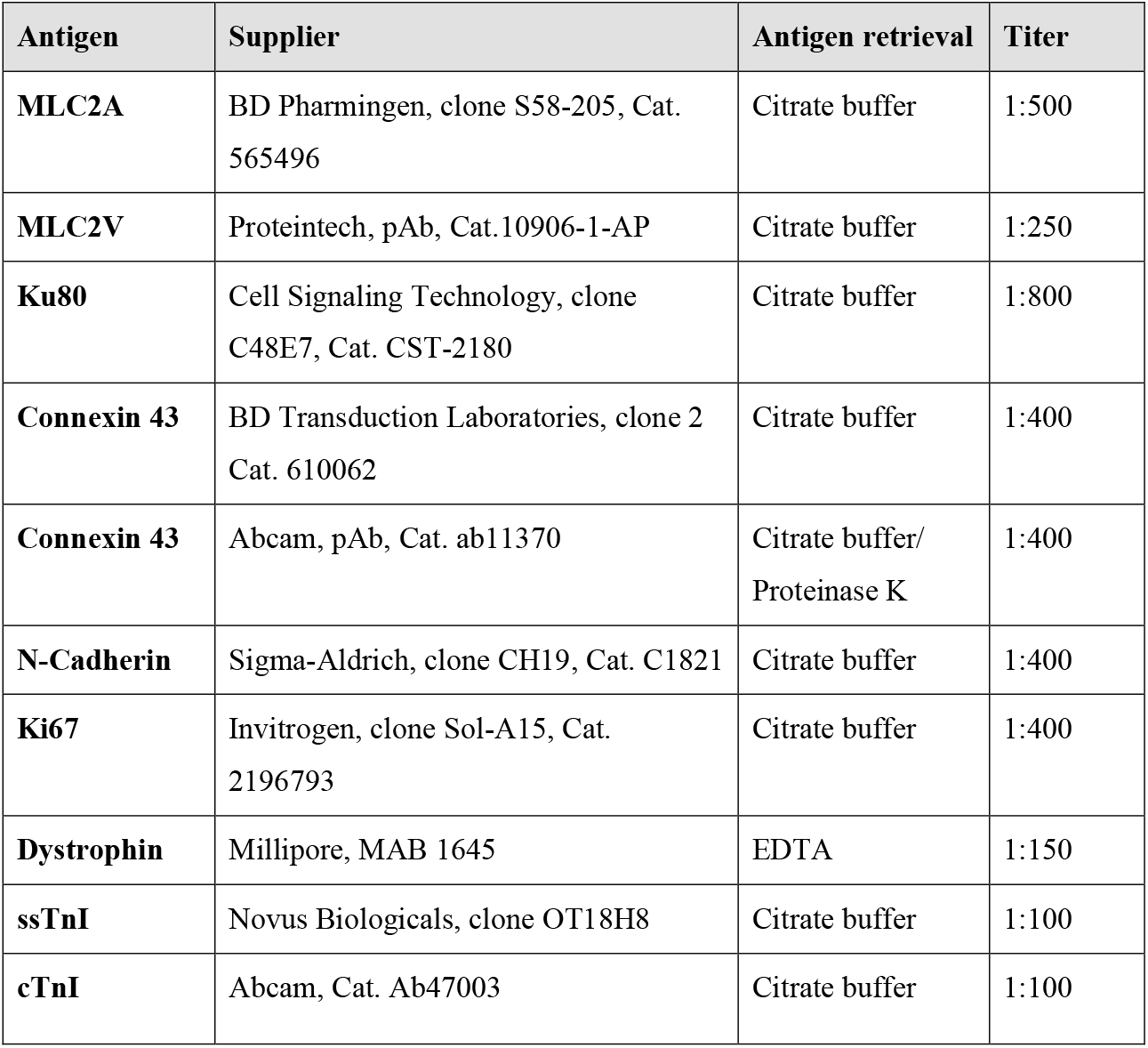
Primary antibodies for histology.

### Langendorff-perfusion

Guinea pigs (body weight 520-700 g) were injected with heparin (1000 U/kg, s.c.) and anesthetized with midazolam (1 mg/kg, s.c.), medetomidine (0,2 mg/kg, s.c.) and fentanyl (0,02 mg/kg, s.c.). Hearts were excised and immersed in ice-cold modified Krebs–Henseleit solution containing (in mM) NaCl 120, KCl 4.7, MgSO_4_ 1.2, NaHCO_3_ 25.0, KH_2_PO_4_ 1.2, glucose 11.1, Na-pyruvate 2.0, and CaCl_2_ 1.8. The aorta was cannulated, and the heart was connected to a custom-made Langendorff apparatus. The heart was perfused using hydrostatic pressure (∼55 mmHg) with warm (36.5±1 °C) modified Krebs-Henseleit solution equilibrated with a mixture of 95% O_2_ and 5% CO_2_ (pH 7.35–7.40). Left ventricular pressure was recorded with a self-assembled balloon catheter, and data were acquired using Chart5 (ADInstruments). After a run-in period of (>10 min), the hearts were illuminated with pulsed stimuli (40 ms pulse duration). Light was applied on the anterior wall of the left ventricle with blue (470 nm, irradiance 1-2 mW/mm^2^) or red light (635 nm, irradiance 1-2 mW/mm^2^) with a pE4000 coolLED for 30-60 seconds. Stepwise increase (0.5 Hz per step) was performed over a time period of 30-45 min.

### Statistics

Statistical analyses were performed with GraphPad Prism 9, USA and R, Vienna, Austria. Comparison among two groups was made by two-tailed unpaired Student’s t-test.

## Author contributions

Tim S. performed and supervised experiments (generation of the BiPOLES iPSC line, cardiac differentiation, animal procedures and Langendorff-experiments), analyzed data, acquired funding and prepared the manuscript. B.S. performed experiments (generation of the BiPOLES iPSC line, cardiac differentiation and Langendorff-experiments), analyzed data and prepared the manuscript. Till S. performed experiments (histology). B.G. conducted animal procedures and performed Langendorff-experiments. C.v.B. conducted animal procedures. J.S.W. provided technical support and conceptual advice for optogenetic experiments. T.E. supervised experiments and prepared the manuscript. F.W. designed the project, supervised and performed experiments (animal procedures, histology, and Langendorff-experiments), acquired funding, and prepared the manuscript.

## Acknowledgments

We thank Kristin Hartmann (UKE, Mouse Pathology Core Facility) for technical assistance in immunohistochemistry. We would like to acknowledge Jutta Starbatty, Thomas Schulze, and Birgit Klampe for technical assistance. We would like to appreciate the contribution of Dr. Sandra Laufer during reprogramming the UKEi001-A line. Flow cytometry was conducted in the FACS Core Facility, UKE. We would like to particularly thank the laboratory animal facility staff (UKE) for their support. This work was supported by a Translational Research Grant from the German Centre for Cardiovascular Research (DZHK; 81X2710153 to TE), the European Research Council (ERC-AG IndivuHeart to TE), and the German Research Foundation (DFG; WE5620/3-1 to FW) and a grant from the Werner-Otto-Foundation (to TS and FW)

## Data Availability

All the data supporting the findings from this study are available within the article and its supplementary information or are available from the corresponding author upon request.

